# Efficient Hydrophilic Small Molecule Delivery To Retinal Cell Lines Using Solid Lipid Nanoparticle Containing Gel Core

**DOI:** 10.1101/2021.05.24.445543

**Authors:** Li Huang, Erico Himawan, Soumaya Belhadj, Raúl Oswaldo Pérez García, François Paquet Durand, Nicolaas Schipper, Matej Buzgo, Aiva Simaite, Valeria Marigo

## Abstract

In this study, we developed a novel solid lipid nanoparticle (SLN) formulation for drug delivery of small hydrophilic cargo to the retina. The new formulation, based on a gel core and composite shell, allowed up to two times increase in the encapsulation efficiency. The type of hydrophobic polyester used in the composite shell mixture affects the particle surface charge, colloidal stability, and cell internalization profile. We validated the SLN capability as a drug delivery system by performing encapsulation of a hydrophilic neuroprotective cyclic guanosine monophosphate analogue, previously demonstrated to hold retinoprotective properties, and the best formulation resulted in particles with size of ± 250nm, anionic charge > -20 mV, and an encapsulation efficiency value of ±60%, criteria that are suitable for retinal delivery. *In vitro* studies using ARPE-19 and 661W retinal cell lines revealed a relatively low SLN toxicity, even when high particle concentration was used. More importantly, SLN could be uptaken by the cells and the release of the hydrophilic cargo in the cytoplasm was visually demonstrated. These findings suggest that the newly developed SLN with a gel core and composite polymer/lipid shell holds all characteristics suitable for drug delivery of small hydrophilic active molecules into retinal cell.

## 1. Introduction

Retinal degeneration is a disease condition characterized by progressive loss of highly differentiated cells within the neurosensory retina, such as photoreceptors, or retinal pigment epithelium (RPE). This condition is commonly found in patients with diabetic retinopathy [1], age-related macular degeneration [2], and hereditary retinal degeneration [3]. Disease associated with retinal degeneration constitutes an important health challenge that severely affects the quality-of-life of patients and has a significant socio-economic impact [4]. Recently, hydrophilic cargos, like nucleic acids [5] or cGMP analogues [6], draw major interest for new development of retinal disease treatments. These hydrophilic cargos may require to be encapsulated in a nanocarrier to reach intracellular targets. However, low encapsulation efficiency, undesired leakage, or initial burst release are common issues occurring during nanoparticulate drug delivery system (DDS) development [7].

Drug delivery to the retina is also notoriously difficult due to the presence of several biological barriers [8,9]. In current clinical practice, intravitreal injections are routinely used to administer treatments to the retina. This route allows DDS to bypass several ocular barriers like the corneal epithelium, the conjunctiva, and the sclera, while the vitreous and internal limiting membrane (ILM) will still need to be crossed. The main considerations of DDS formulation in intravitreal injections routes are the particle size, surface charge, and the functional materials used to enhance intracellular delivery. The vitreous has a gel-like network structure with a pore size expected to be in the range of 500 nm [10]. Small drugs, proteins, and nanoparticles may diffuse through the vitreous [10,11]. A negatively charged surface and size below 500 nm are preferred qualities for DDS because these features may improve particle mobility [11,12]. When the drug requires intracellular targeting within the retina, barriers at the level of the plasma membrane will limit its diffusion, especially for hydrophilic molecules [13]. A DDS can help the delivery of the drug [8] and nanoparticles composed with material that enhances endocytosis, such as solid lipid nanoparticles (SLN), will likely improve the entrance of the active compound into the target cells [9,14,15].

The water-in oil-in water (W_1_/O/W_2_) emulsion method is generally used to prepare lipid-based particles to encapsulate small hydrophilic molecules. Lipid-based particles obtained via this method, such as liposomes, often have a low encapsulation efficiency due to leakage from its core during preparation [16]. Other lipid-based particles, like conventional SLN, are limited in space for uploading hydrophilic molecules [17] and compounds may be expelled following the polymorphic transition of the structure during storage [18]. To accommodate larger amounts of active molecules in the core of lipid-based particles, and prevent their premature release, the addition of a thermo-responsive gel core [19] or of micelles [20,21] was shown to improve the encapsulation of the hydrophilic cargo. Unfortunately, these studies used a large macromolecule (*i*.*e*. protein) as cargo and direct translation of their DDS suitability for delivering small hydrophilic molecules into retinal cells is not readily available.

In this study, we evaluated the potential of adapting and modifying SLN formulations, originally intended for holding hydrophilic macromolecules, as a suitable DDS for delivering small hydrophilic molecules into retinal cells. Firstly, we investigated the effect of adding a thermo-responsive gel core inside the SLN in terms of encapsulation efficiency. We also investigated the effect of adding hydrophobic polyester to the SLN shell in regard to DDS size, surface charge and polydispersity. The small size, negative surface charge, stability, encapsulation efficiency, low toxicity, and internalization capability in retinal cells *in vitro* support the possible use of such SLN for intravitreal delivery.

## 2. Materials and Methods

### 2.1 Materials

Poloxamer 407 (Sigma Aldrich), poloxamer 188 (Applichem), rhodamine B (RhoB; Sigma Aldrich), Rp-8-Br-PET-cGMPS, also known as CN03 (provided by Research Institute of Sweden), Glycerol tripalmitate (GTP; Alfa Aesar), soy-bean lecithin (LCT; VWR), stearic acid (SA; BASF), 50/50 DL-lactide/glycolide copolymer (PLGA; Corbion), poly-ε-caprolactone 14kDa (PCL; Sigma Aldrich), 8-aminonaphthalene-1,3,6-trisulfonic acid disodium salt (ANTS; Biotium), p-Xylene-Bis-Pyridinium Bromide (DPX; Biotium), dichloromethane (VWR), deionized water (VWR), Human adult retinal pigment epithelial cells (ARPE-19 cell, ATCC), 661W cell (generously provided by Dr. Muayyad Al-Ubaidi, University of Oklahoma), Dulbecco’s modified Eagle’s medium and Ham’s F12 nutrient mixture (DMEM/F12; Gibco), low glucose (1mg/ml) Dulbecco’s modified Eagle’s medium (DMEM; Gibco), fetal bovine serum (FBS; Gibco), glutamine (Sigma Aldrich), penicillinstreptomycin (Sigma Aldrich), Accutase® solution (Sigma-Aldrich), paraformaldehyde (PFA; Sigma Aldrich), anti-Zonula occludens-1 (ZO-1) antibody (Invitrogen), goat anti-rabbit secondary antibody (Life technologies), 4’,6-diamidino-2-phenylindole, dihydrochloride (DAPI; Sigma Aldrich), colorimetric methyl-thiazolyl diphenyl-tetrazolium bromide (MTT; Sigma Aldrich). All purchased materials were used as received.

### 2.2 Nanoparticle Synthesis

The SLN formulation is illustrated in Figure 1A. A stock solution for W_1_-phase without gel core was prepared by dissolving RhoB in deionized water to reach 10 mg/ml concentration. For W_1_-phase stock with gel core, poloxamer 407 was added to the RhoB solution to reach 40% w/v. To ensure complete dissolution, the poloxamer 407 solution was dissolved at 4°C for 48 h. The W_2_*-*phase stock solution was prepared by dissolving poloxamer 188 in deionized water to reach 2% w/v concentration. The O-phase solution was prepared by dissolving GTP, LCT, SA, and PCL or PLGA in 1 ml of dichloromethane according to different formulation codes listed in Supplemental Table 1. Finally, for the preparation of blank and drug-loaded particles, deionized water or CN03 were added respectively in the W_1_-phase instead of RhoB. Similarly, for release assays of the compound from SLN inside the cell, ANTS (25 μM) and DPX (90 μM) were co-encapsulated in the W_1_-phase instead of RhoB. ANTS and DPX, as a pair of fluorescence tracer and quencher, were chosen based on previously published studies [22].

**Figure 1.**
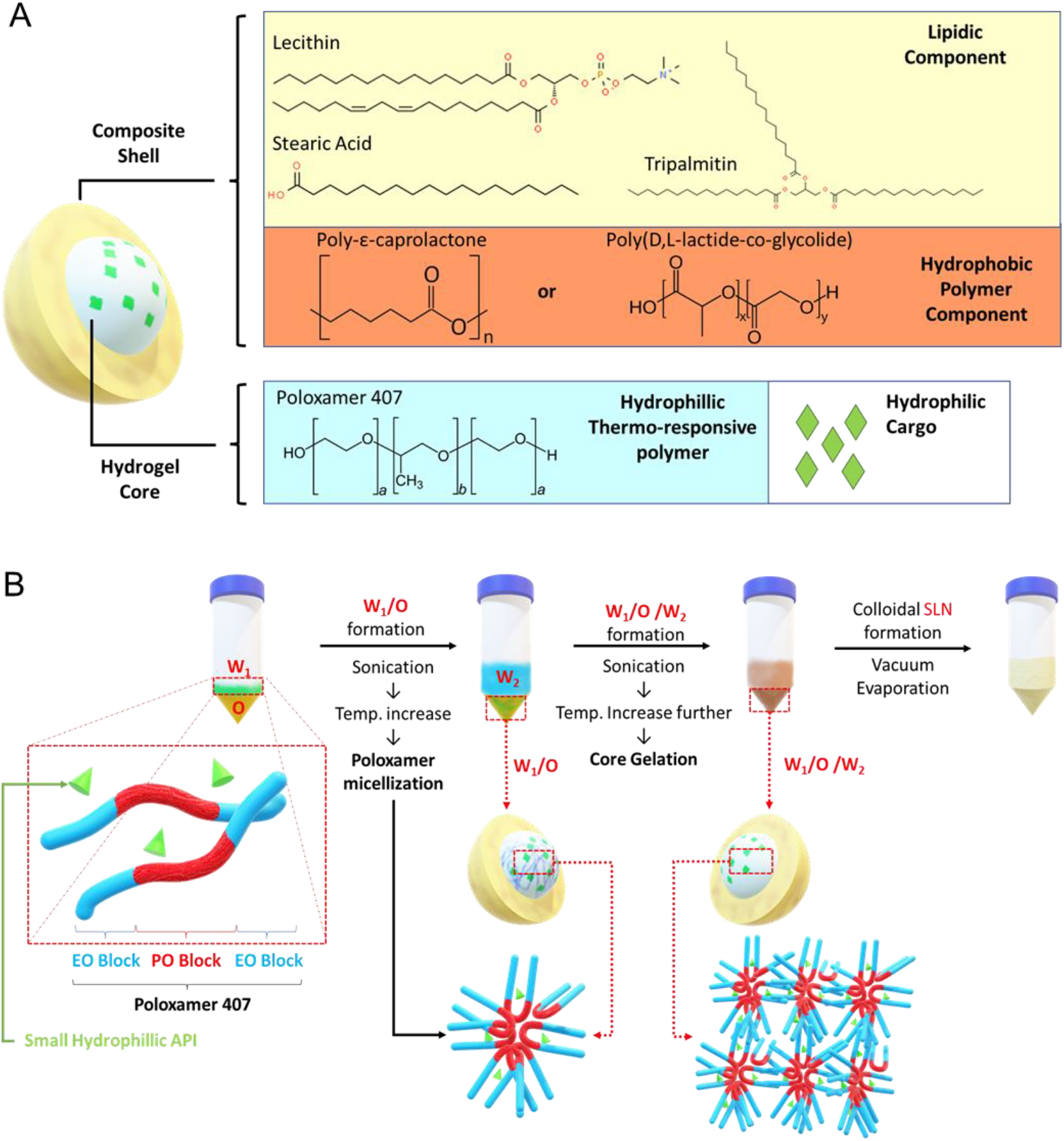
Solid Lipid Nanoparticle (SLN) Formulation. (A) Schematic representation of SLN containing gel core and composite shell. (B) Core gelation procedure of thermo-responsive poloxamer 407.

The synthesis was performed by adding 200 μl of W_1_-phase stock solution kept at 4- 7°C, using an ice bath, to 1 ml of O-phase solution followed by sonication with the Q55 ultrasound probe from Qsonica (amplitude: 30% for 60 s without a pulse) to form the primary W_1_/O emulsion. Then, 4.8 ml of W_2_-phase solution was added into the primary emulsion and sonicated to form W_1_/O/W_2_ emulsion (amplitude: 40% for 10 s followed by 20 s at 20% amplitude without a pulse). The solution was further diluted with 10 ml of W_2_- phase solution followed by sonication (amplitude: 20% for 30 s without a pulse). The organic solvent was then removed by vacuum evaporation (P: 650 mmHg) at room temperature for 20 minutes to form the nanoparticles. The resulting colloidal solution was stirred for 4 h to ensure complete removal of dichloromethane.

### 2.3 Physicochemical characterization

#### Dynamic laser scattering

The particle size distribution and hydrodynamic diameter were measured using NanoPhox DLS equipment from Sympatec, GmbH. Before the measurement, concentrated particle solutions were diluted 5 times using 2% w/v poloxamer 188 solutions. The analyses were performed using a non-negative least square (NNLS) algorithm integrated in Windox5 software from Sympatec, GmbH. Viscosity of the solutions was calibrated and validated using the polystyrene bead standard. For each sample, the measurement was repeated 3 times and each lasted 200 seconds.

#### Zeta potential

The zeta potential of the particles was measured by Zetasizer equipment from Malvern. Samples were diluted 5 times using 10 mM phosphate buffer, pH 7.4. For each sample, the measurement was repeated 3 times.

#### Morphological analysis

SLN solutions, as described in supplemental table 1, were synthesized without hydrophilic cargo (*i*.*e*. blank particle) and analyzed using a transmission electron microscope (TEM). The colloidal nanoparticle solutions were stained with 7% (w/v) phosphotungstic acid as negative contrast. The morphological characterization of the particle was performed at a TEM acceleration voltage of 120 kV.

#### Encapsulation Efficiency

Encapsulation efficiency (EE) was measured using an indirect method in which the amount of unencapsulated cargo outside the particles was measured. Sample solutions were filtered using a 100 kDa microcentrifuge membrane filter (Sartorius) (3 × 5 minutes, @5000 RCF). The filtrate was collected, and the amount of cargo was quantified using equation 1. As there might be some loss of W1-Phase in the pipette tips during synthesis, cargo loss was quantified to avoid encapsulation efficiency overestimation.

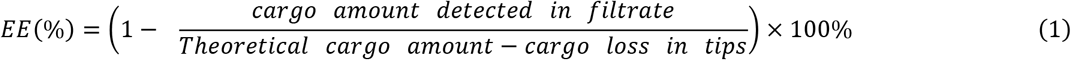

RhoB and CN03 concentrations were quantified by measuring absorbance at 550 nm and at 254 nm using an UV-spectrophotometer (Biomolecular device) and high-pressure liquid chromatography (HPLC) system (Dionex Ultimate 3000), respectively. For HPLC, mobile phases A and B were 5 mM ammonium acetate buffer, and acetonitrile, respectively. 5μl of the sample was injected into the column (Waters^®^ XBridge C18 XP column, 50 × 3 mm). The HPLC quantification was performed using the Chromeleon software by calculating the peak area in the retention time of around 3.1 minutes. The experiment was replicated 3 times for both UV-spectroscopy and HPLC.

#### Stability study

Colloidal stability was assessed by monitoring changes in size for the resulting nanoparticle over time. Concentrated SLN solution was diluted in 10 mM PBS to reach 200 μg/ml concentration and stored in glass vials at either 25°C or 4°C, for a period of 4 weeks. At each time point, the vials were gently shaken by hand, brought to room temperature prior and analyzed by Dynamic laser scattering.

### 2.4 Cell cultures

The cells primarily affected in retinal degeneration are RPE and photoreceptors. For this study we chose two retinal cell types: i) ARPE-19, a spontaneously arising human RPE cell line with normal karyotype [23]; ii) 661W photoreceptor-like cells, derived from a mouse retinal tumor generated in a transgenic mouse expressing the SV40 large T-antigen under the control of the *IRBP* (interphotoreceptor retinoid-binding protein) promoter [24]. ARPE-19 cells were cultured in DMEM/F12 supplemented with 10% FBS and 1% penicillin-streptomycin in an incubator at 5% CO_2_ and 37°C. The murine photoreceptor 661W cell line was cultured in low glucose (1 mg/ml) DMEM supplemented with 10% FBS, 2mM glutamine and 1% penicillin-streptomycin in an incubator at 5% CO_2_ and 37°C. Approximately every three days cells reached 70–80% confluence and were sub-cultured.

### 2.5 Fluorescence microscopic analysis and immunofluorescence

Cells were seeded on glass coverslips in a 24-well plate at a density of 4× 10^4^ cells/well. 24 h later, cells were exposed to RhoB loaded SLN (RhoB-SLN). As control, cells were treated with free RhoB solution at the same concentration (20 μM) of RhoB present in the nanoparticle solution. A blank was prepared by incubating the cells with nanoparticles containing no fluorophore.

For compound release assays, 24 h after seeding, ARPE-19 cells were treated with medium containing 200 μg/ml blank SLN (blank); freely dissolved tracer (ANTS); free tracer together with quencher (free ANTS/DPX), or 200 μg/ml SLN loaded with ANTS/DPX (ANTS/DPX-SLN). 5 hours later the medium was replaced with fresh medium without any particles or fluorophores and the incubation was continued for 24 h, 48 h or 72 h.

After incubation, cells were rinsed with phosphate buffer saline (PBS), fixed with 2% PFA for 10 minutes and nuclei were stained with 0.1 μg/ml DAPI. For the immunofluorescence, cells after fixing were incubated with anti-ZO-1 (1:100) primary antibody overnight at 4°C. After washing three times with PBS, cells were incubated with the Alexa Fluor^®^ 488 goat anti-rabbit secondary antibody (1:1000) and 0.1 μg/ml DAPI for 40 min at room temperature. Slides were mounted with Mowiol 4–88 and cells were observed using the Zeiss Axio Imager A2 fluorescence microscope. Mean Fluorescence Intensity (MFI) of each cell was quantified by ImageJ software (n_cells_ ≥ 10).

### 2.6 Cell Viability Assay

Cell viability assay was performed by colorimetric methyl-thiazolyl diphenyl-tetrazolium bromide (MTT) assay. Cells were seeded on 96-well plates at a density of 6,000 cells/well. After treatment with SLN for various times, the medium was aspirated, and cells were incubated with 50 μl of 1 mg/ml MTT solution for 90 minutes at 37°C. The supernatant was removed, and the purple formazan crystals were dissolved in 100 μl isopropanol. The plate was shaken for 10 minutes and analyzed at 570 nm using a microplate reader (Labsystems Multiskan MCC/340).

### 2.7 Flow cytometry analysis

ARPE-19 and 661W cells were seeded on 12-well plates at a density of 1 × 10^5^ cells/well. After treatment with control or SLN, cells were detached with 500 μl Accutase^®^ and collected by centrifugation at 300xg for 5 minutes at room temperature. The cells were washed three times with 500 μl PBS and collected by centrifugation at 300xg for 5 minutes at room temperature. The cell pellet was resuspended with 500 μl of PBS and RhoB fluorescence was immediately analyzed at the Attune^®^ NxT Acoustic Focusing Cytometer. The channel voltage and gain were maintained constant throughout the whole analysis.

### 2.8 Statistical analysis

Data are presented as the means ± SEM. Student’s t tests were applied to compare two groups. Analysis of variance (ANOVA) was used for comparisons of data with greater than two groups. Post hoc comparisons were performed with Bonferroni test. Significance was set at *P<0.05, **P<0.01, ***P<0.001. All statistical analyses were performed using SPSS (Statistics 21; IBM Inc., Bentoville, AR, USA).

## 3. Results

### 3.1 Generation and characterization of solid lipid nanoparticles containing a gel core

The aim of this study was to develop a drug delivery system (DDS) to facilitate uptake of hydrophilic molecules by retinal cells. Among common thermo-responsive gels, such as poloxamer 407, chitosan, and hydroxypropyl methylcellulose (HPMG) which have been previously studied [19], we chose poloxamer 407, as the gel core material, because gelling can be easily induced by increasing temperature. For the lipidic shell, we used the mixture of lecithin (LCT), tripalmitin (GTP), and stearic acid (SA), since this mixture has been previously reported to be able to significantly enhance nanoparticle cellular uptake [25]. In addition, we added hydrophobic polyester to the lipid mixture to create a composite shell using biocompatible PCL and PLGA (Figure 1A). To allow evaluation of encapsulation and cellular uptake, we chose a small hydrophilic cargo Rhodamine B (RhoB; 479.02 g/mol) that can be easily tracked during the experiments. The illustration of the procedure for SLN generation with a size range of 200-250 nm containing a gel core can be seen in Figure 1B.

The summary of the generated SLN components is reported in Supplementary Table 1. The addition of hydrophobic polyester such as PCL and PLGA may improve particle polydispersity index (PDI) to less than 0.4 when used in combination with a gel core. Moreover, all of the produced nanoparticles were anionic as characterized by their zeta potential. The addition of gel core didn’t significantly affect the surface charge, *e.g*., SLN.03 (−27± 2.3 mV) versus SLN.06 (−24± 1.5 mV). On the other hand, the presence of the gel core improved the encapsulation efficiency, *e.g*., SLN.02 (24± 0.8%) versus SLN.05 (48 ± 0.44%). Both types of particles with a gel core could encapsulate Rhodamine B above 40%, while particles with an aqueous core had RhoB encapsulation efficiency at around 20%, regardless of the shell type. The addition of hydrophobic polyester to the shell formulation had no detectable effect on the encapsulation efficiency (Figure 2A). The SLN morphology, observed using TEM, confirmed that all the produced particles were far below 500 nm, fulfilling the basic size requirement for mobility in the vitreous (Figure 2B).

**Figure 2.**
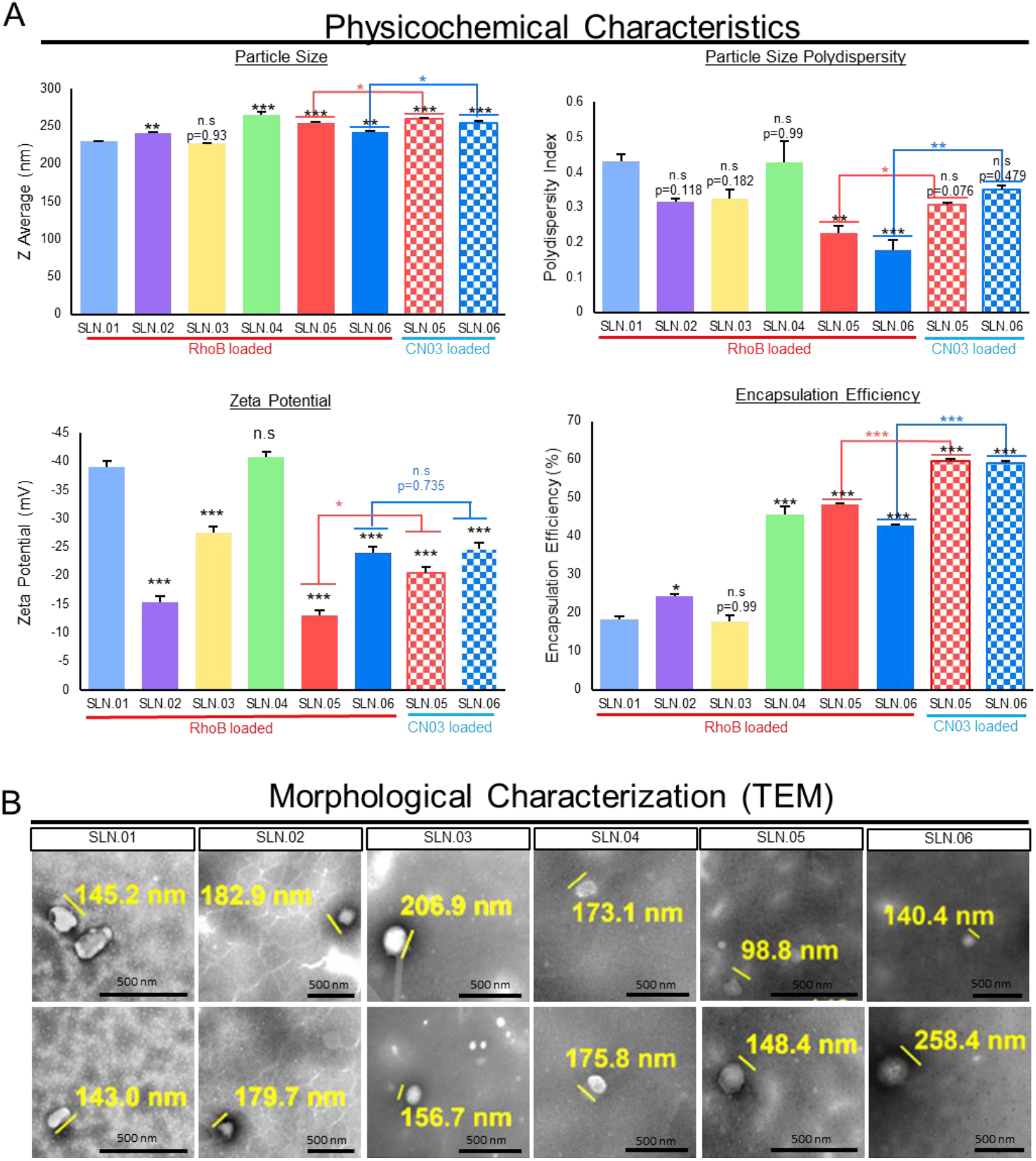
Characterization of SLN. (A) Physicochemical characteristics of the generated SLN compared to conventional SLN (SLN. 01). Data are presented as mean ± SEM, n=3 independent experiments. Significance at *P<0.05, **P<0.01, ***P<0.001, ANOVA followed by Bonferroni’s post hoc test. Statistical analysis comparing SLN.05 and SLN.06 loaded with either RhoB or CN.03 were performed using Student’s t test. (B) Representative image of morphological analysis of blank SLN without cargo using TEM. Scale bar: 500 nm.

Based on the analytical analyses of the generated SLN, we chose to focus on SLN.05 and SLN.06 for further studies since they showed an improvement in PDI and encapsulation efficiency compared to conventional SLN with pure lipid shell and no gel core (SLN.01). We firstly validated the formulation encapsulation capability of SLN.05 and SLN.06 using a known compound previously shown to have neuroprotective properties in the retina, the compound called CN03 [26]. Both SLNs have been able to encapsulate CN03 and resulted in negatively charged particles in the similar range of 200-250 nm. We noticed an even better encapsulation efficiency (i.e., ±15% increase) compared to RhoB when CN03 was used (Figure 2A). A colloidal stability study was then performed with CN03-loaded SLN dispersed in a PBS solution. The samples were stored at different storage temperatures for 4 weeks. Increase in size (±30 nm) was observed in the SLN.05 colloidal solution after one week of storage (data not shown). In comparison, SLN.06 colloidal solution showed better size stability within the study period. More importantly, the particle size was maintained below 300 nm regardless of storage temperature within 1 month of storage (Figure 3).

**Figure 3.**
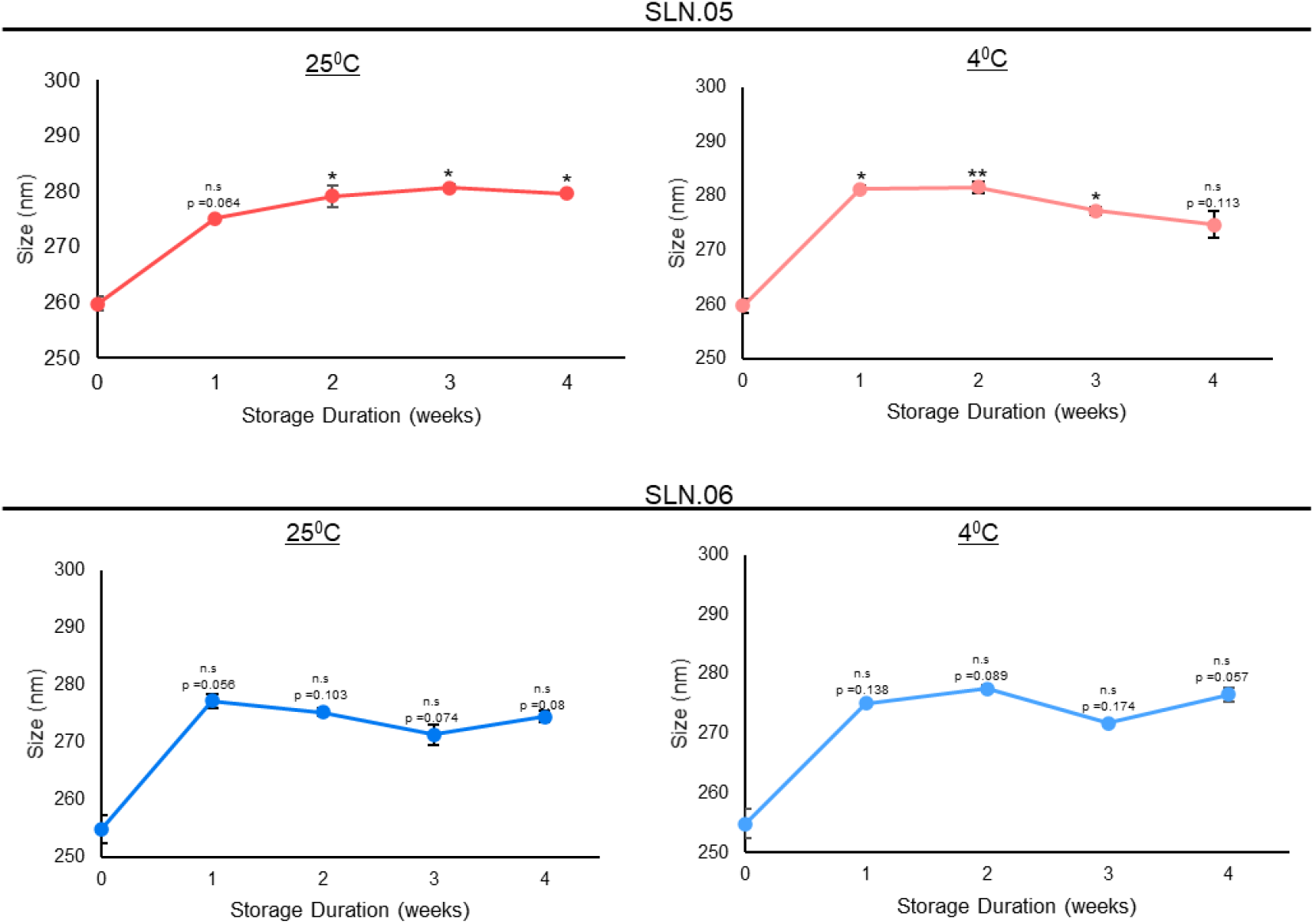
Stability study of nanoparticle containing CN03 in 10 mM PBS at different storage temperature. Stability of SLN.05 and SLN.06 stored at 25°C and 4°C was evaluated by analyzing the particle sizes every week up to 4 weeks in PBS. Data are presented as mean ± SEM, n=3 independent experiments. Significance at *P<0.05, **P<0.01, ***P<0.001, ANOVA followed by Bonferroni’s post hoc test.

Taken together, all physicochemical characterization data show that SLN.05 and SLN.06 have a relatively good property to be further *in vitro* validated as a potential DDS for retinal cells in this study.

### 3.2 Evaluation of SLN.05 and SLN.06 toxicity to ARPE-19 and 661W retinal cell lines

We used ARPE-19 cells (human retinal pigment epithelium cell line) and 661W cells (mouse photoreceptor-like cell line) to evaluate the toxicity of SLNs on retinal cells. We exposed ARPE-19 and 661W cells to either SLN.05 or SLN.06 at different concentrations and evaluated toxicity by the MTT cell viability assay at different time points.

Both SLN.05 and SLN.06 showed increase toxicity in dose-dependent manner and time-dependent manner (Figure 4A-B). Overall, SLN.05 showed higher toxicity in both cell types. Toxicity of SLN.05 on ARPE-19 cells started to be detected at 200 μg/mL after 5 hours of exposure. Interestingly, 661W cells showed higher resistance to SLN.05 toxicity because 200 μg/mL of SLN.05 did not significantly reduce 661W viability even after 24 hours of exposure but started to be toxic at 500 μg/mL. SLN.06 demonstrated to be less toxic to both cell lines, especially to 661W cells. ARPE-19 cells could tolerate up to 500 μg/mL of SLN.06 within 5 hours of exposure, while 661W cells could tolerate up to 800 μg/mL of SLN.06 within 24 hours exposure. SLNs reduced viability of both ARPE-19 and 661W cells after 48 hours of exposure (Figure 4A-B).

**Figure 4.**
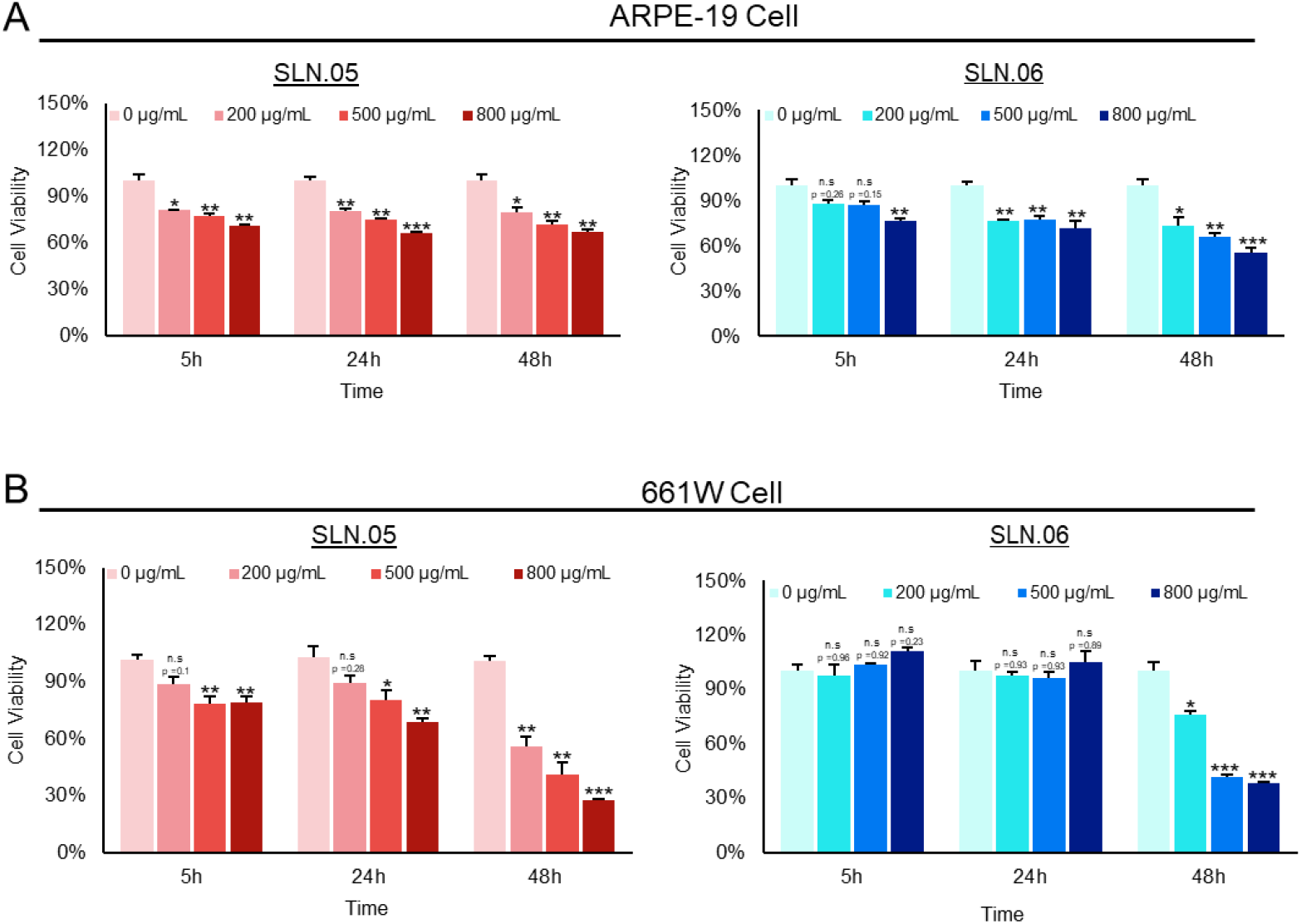
Toxicity of SLN to ARPE-19 and 661W retinal cell lines. Toxicity of SLN.05 and SLN.06 to retinal cells line was assessed by measuring the percentage of cell viability after exposure of SLN at various concentrations and at different time points using MTT assay. (A) Percentage of ARPE-19 cells viability after exposure of SLN.05 or SLN.06. (B) Percentage of 661W cells viability after exposure of SLN.05 or SLN.06. Untreated cells (0 μg/mL) were set as 100% cell viability and used as a control. Data are presented as mean ± SEM, n=3 independent experiments. Significance at *P<0.05, **P<0.01, ***P<0.001, ANOVA followed by Bonferroni’s post hoc test.

### 3.3 SLN.05 and SLN.06 internalization by retinal cell lines

To deliver small hydrophilic molecules to retinal cells, SLN need to be efficiently internalized by cells. To visualize SLN uptake, we used Rhodamine (RhoB) loaded SLN (RhoB-SLN). Both RhoB-SLN.05 and RhoB-SLN.06 could be efficiently internalized by 661W cells, where RhoB intensity in the cytosol increased in a concentration dependent manner (Figure 5A-B). To further confirm and quantify internalization efficiency of SLNs, we exposed 661W cells to 200 μg/mL of RhoB-SLN.05 and RhoB-SLN.06 and quantified the fluorescence signal at different time points by flow cytometry. Since free RhoB also can penetrate the cells, we used cells treated with free RhoB suspension for 5 hours as control. Only 0.08% of 661W cells were positive for RhoB after treated with free RhoB for 5 hours, indicating that RhoB diffusion inside the cells was very low. 1 hour exposure to RhoB-loaded SLN was sufficient to detect 3.28% of 661W cells positive RhoB after incubation with RhoB-SLN.05 and 3.35% of 661W cells positive after incubation with RhoB-SLN.06. The percentage of RhoB positive cells increased with longer incubation times (Figure 5C). Similarly, we observed the same trend of SLN uptake with ARPE-19 cells (Supplemental figure 1). Taken together, these data indicate that SLN.05 and SLN.06 can be internalized by both photoreceptor and RPE cell types.

**Figure 5.**
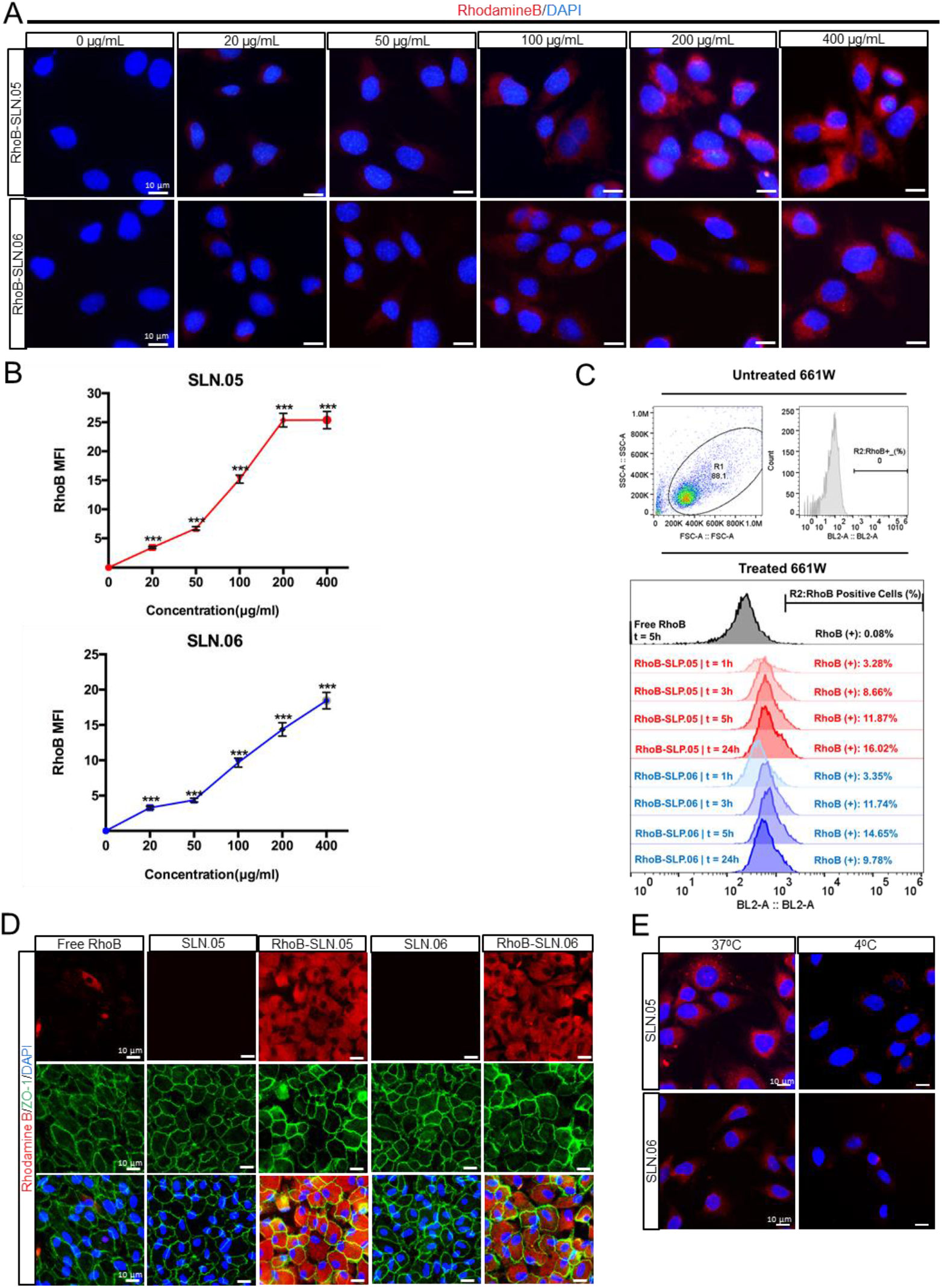
Internalization of SLN.05 or SLN.06 by retinal cell lines. Visualization of internalized SLN by 661W cells using Rhodamine B-loaded SLN (RhoB-SLN). (A) Micrographs of internalized RhoB-SLN.05 or RhoB-SLN.06 (red signal) at various concentrations after 5 hours of exposure. Nuclei of cells were stained with DAPI in blue. Scale bar, 10μm. (B) Mean fluorescence intensity (MFI) of Rhodamine B signal inside the cells were quantified using image J software (n_cells_ ≥ 10). (C) Histogram overlay of RhoB relative fluorescence intensity and percentage of RhoB positive cell (RhoB+) assessed by Flow cytometry. R1: gating for selecting cells population; R2: gating to select RhoB+ cells based on blue laser (BL2-A) detector; FSC-A: forward scattering channel; SSC-A: side scattering channel. (D) Micrographs showing intracellular localization of RhoB-SLN.05 or RhoB-SLN.06 (red signal) in ARPE-19 cells. Cell membranes were stained with anti-ZO-1 antibody (green). Nuclei of cells were stained with DAPI in blue. Scale bar, 10μm. (E) Representative images showing temperature-dependent RhoB-SLN.05 or RhoB-SLN.06 (red signal) uptaken by 661W cells at either 37°C or 4°C. Nuclei of cells were stained with DAPI in blue. Scale bar, 10μm.

We visually confirmed the intracellular localization of SLNs after being internalized by the ARPE-19 cells by staining the membrane of the cells with an anti-ZO-1 antibody (specific antibody that recognize a peripheral membrane protein in epithelial cells), and we observed that RhoB-SLNs are localized inside cytosol after the internalization process. (Figure 5D). To elucidate the mechanism of internalization of SLN.05 and SLN.06 by the photoreceptor cells, we exposed 661W cells to 200 μg/mL of RhoB-SLN.05 and RhoB-SLN.06 and incubated for 1 hours at either 37°C or 4°C. We observed that incubation at 4°C highly limited the uptake of Rho-SLNs, indicating an energy-dependent process rather than passive membrane passage (Figure 5E). Based on the knowledge that most of nanoparticles are internalized by cells through endocytosis [27], these data confirmed that SLN.05 and SLN.06 were uptaken via the endocytic process rather than membrane permeation.

### 3.4 Encapsulated cargo release inside the cells

To evaluate if SLN.05 and SLN.06 can successfully release the cargo after being taken up by the cells, we performed fluorescence leakage assay using ANTS/DPX which has been widely used to study vesicle leakage [28]. We encapsulated the ANTS fluorescent dye together with its quencher DPX. Once the SLN shell breaks and releases the cargo inside the cells, DPX will not be able to quench ANTS anymore due to the increase of the molecular distance between ANTS and DPX, which leads to low collision probability and allowing free ANTS inside the cells to emit green fluorescence (Figure 6A). In this experiment we exposed cells to either free ANTS or DPX, that are not able to penetrate the cells, as control. Only cells exposed to SLN.05 and SLN.06 loaded with ANTS/DPX resulted fluorescent demonstrating that SLNs could successfully deliver ANTS/DPX inside the cells and release the cargo (Figure 6B-C). A faint signal could be detected at 24 hours, but a full signal was easily detected after 48 hours of exposure (Figure 6C). Taken together, these data demonstrate that the new formulated SLNs are able to release a hydrophilic molecule inside a retinal cell and can be an efficient drug delivery system for the retina.

**Figure 6.**
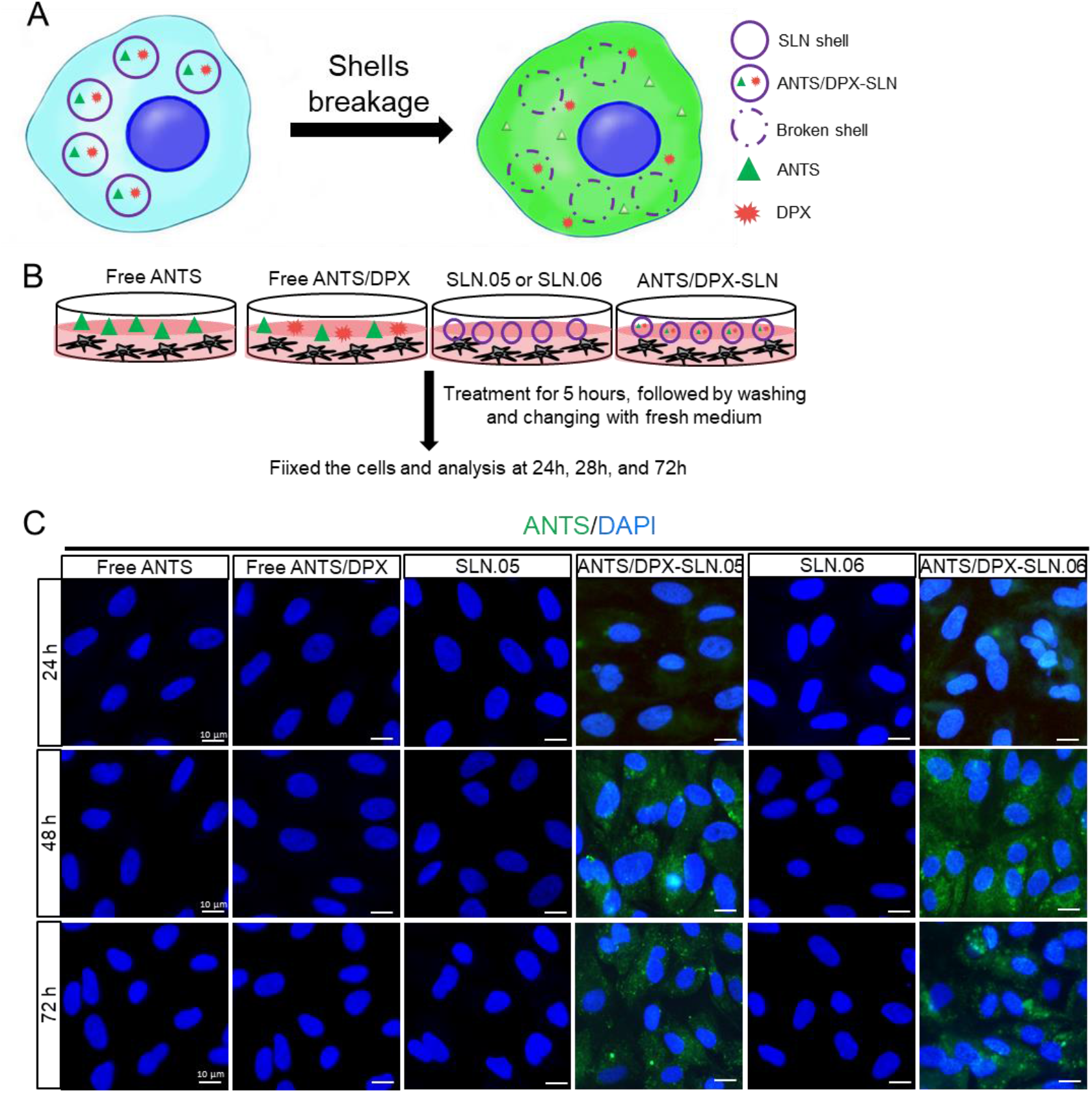
SLN.05 or SLN.06 cargo release inside the cells. Fluorescence leakage assay using ANTS/DPX was used to determine cargo delivery and release by the SLN.05 or SLN.06 inside the cells. (A) Schematic summary of fluorescence leakage assay using ANTS/DPX. (B) Schematic of experimental design for fluorescence leakage assay. (C) Micrographs of ARPE-19 cells exposed to ANTS/DPX loaded SLN.05 and SLN.06. Released ANTS was detectable in green only in cells exposed to ANTS/DPX loaded into SLNs but not in cells exposed to free ANTS and/or DPX. Nuclei of cells were stained with DAPI in blue. Scale bar, 10μm.

## 4. Discussion

The delivery of a drug to the neural retina is challenging due to the different barriers that need to be crossed and the physicochemical environment of the vitreous that may affect the passage of the drug to the target cells. In this study we presented new formulations of nanoparticles which can enter retinal cells while having features that may facilitate navigation across the vitreous (e.g., size < 500 nm and anionic). The key findings from the formulation development studies were: (*i*) the gel core improved the encapsulation efficiency by up to 2-fold; (*ii*) the addition of hydrophobic polymer to the shell could be used to tailor the surface charge of the final DDS. Our results on encapsulation efficiencies suggested that the gel core could improve the small hydrophilic cargo retention during formulation. This agrees with previous reports that used large macromolecules as cargo [19]. Most likely, the improved retention came from the solidification of Poloxamer 407 emulsion droplets. Poloxamer 407 droplets formed a nanogel thanks to a local increase in temperature during the sonication process. Particle surface charge should be considered in terms of cellular uptake. Cellular membrane is generally negatively charged and thus, a strongly anionic particle will more difficultly enters the cells compared to a cationic particle [29]. When the particle is cationic however, it will have tendency to aggregate in vitreous [11]. Thus, there is a need to tailor particle surface charge during DDS development.The addition of polyester to the shell formulation reduced the strong negative charge of the pure lipid SLN shell (SLN.01; -39 mV). The intensity of the surface charge reduction differed based on the hydrophobic polymer used as a filler in the composite SLN shell formulation (i.e., PCL and PLGA were used in this study). Particle shells containing PCL (SLN.02; -15 mV) had a higher zeta potential reduction compared to PLGA (SLN.03; -27 mV). The charge reduction, observed from adding PCL or PLGA to create a composite shell, may indicate that the hydrophobic polymeric chains are well distributed on the surface. The intensity of surface charge observed is very likely related to the inherent surface charge of the polymer used. Based on this finding, the choice of hydrophobic polymeric components in the composite SLN shell may be used to tailor specific surface charges in further stage of DDS development.

The SLN formulation initially developed with RhoB as the hydrophilic cargo was validated with a real drug for retinal degeneration (i.e., CN03). The freshly synthesized CN03-loaded SLN particle size was maintained in the range of 200-250 nm. There was a significant change in surface charge when CN03 salts were used instead of RhoB for SLN.05. Yet, this was not observed in SLN.06. Without CN03 salts, SLN.05 (−13 mV) had a less negative charge compared to SLN.06 (−24 mV). Thus, unencapsulated CN03 salts had a weaker influence or had a less surface absorption on a more negatively charged SLN. We also observed an increase in polydispersity when CN03 salts were used as cargo. Colloidal system is a delicate particulate system which is strongly influenced by the salts and pH from the dispersing medium. The increase in polydispersity might come from effect of unencapsulated CN03 salts during the synthesis. Finally, there was about 15% increase in encapsulation efficiency of CN03 compared to RhoB. This may be due to the fact that CN03, which is in a sodium salt form, has a much lower solubility in dichloromethane compared to RhoB. RhoB, thus, can more easily leaks out from the W_1_ phase during the DDS preparation compared to CN03. Based on this finding, we surmise that this DDS may also work for other hydrophilic cargos with lower solubility in the organic solvent (*e.g*., DNA) for different pharmaceutical application.

The colloidal stability study, which was the last checkpoint in this work prior to *in vitro* study, showed a ±30 nm increase in particle size for SLN.05 after the first week of storage. The observed increase in particle size may be caused by the high concentration of salts from PBS disrupting the colloidal solution stability. However, SLN.06, which had a more negatively charged surface compared to SLN.05, showed a statistically better stability profile throughout the study. This observation might come from the fact that the magnitude of particle-to-particle repulsion, which could help prevent aggregation, is proportional to the intensity of the particles surface charge. While SLN.06 may seem to perform better than SLN.05 in terms of prolonged colloidal stability in salt solution, both SLN size was maintained below 300 nm throughout the stability study regardless of the storage temperature and duration. Thus, both SLN.05 and SLN.06 were selected for *in vitro* study using 661W and ARPE-19 cells.

While it was clear that both SLN.05 and SLN.06 showed a time and dose dependent toxicity towards the cells, the *in vitro* cytotoxicity studies provided the insight that the two retinal cell lines had different sensitivity to these SLNs. This agrees with previous studies reporting that distinctive cell physiology, proliferation rate, metabolic activity, membrane, and phagocytosis characteristics are responsible for the different sensitivity to external factors [29,30]. Physicochemical elements of nanoparticles can also affect the cytotoxicity of cells [29]. Specifically, distinct shell composition of SLN.05 and SLN.06 may differently affect viability of retinal cells. With regards to cytotoxicity, SLN.06 seemed to perform better than SLN.05 as a DDS.

Internalization studies in two retinal cell types demonstrated that: (*i*) the SLN formulation helped internalization of small hydrophilic compounds; (*ii*) the SLN shell component might be used to tailor the uptake rate in different cell types. We observed that ARPE-19 cells could better uptake SLN containing PCL in the shell (SLN.05). This might be due to the fact that SLN.05 is less negatively charged compared to SLN.06 (Figure 2A) and the uptake level is directly affected by physicochemical properties of SLN, such as shape, size and surface charge [31]. In 661W cells, the uptake profile of SLN.06 nanoparticle was similar to the one measured in APRE-19 cells and limited to a low percentage of cells that internalized the nanoparticles. For SLN.05, less uptake was observed in 661W compared to ARPE-19. This difference may be attributed to the fact that uptake rates are also specific to each cell type [32]. It is not surprising that photoreceptor cells have a lower uptake rate compared to ARPE-19 cells, because RPE cells have a high rate of phagocytosis, in fact one of their functions is to daily phagocytose photoreceptor outer segments [33]. The reduced uptake at 4°C suggested that the SLN mainly enter the cells via endocytosis, as the energy dependent endocytosis will be largely inhibited at this temperature [34–36]. We also demonstrated that SLN could release their cargo after being internalized by the cells. This result highlights that the newly developed DDS can be appropriate for encapsulation of small hydrophilic drugs and for their release into the target cells.

Overall, both SLN.05 and SLN.06 can successfully improve the uptake of small hydrophilic cargo into retinal cell lines *in vitro*. However, SLN.06 seems to perform better as a DDS when compared to SLN.05 considering its slight advantage in terms of stability and cytotoxicity. Finally, while the relatively simple cell culture environment yielded interesting data, a full drug/DDS efficacy testing will likely require more complex test systems. More advanced *in vitro* tests using organotypic retinal explant cultures, in which the normal histotypic context of the retina is preserved [37], or *in vivo* injections will further characterize the suitability of the new SLN for the delivery to the retina. The fate of SLN materials after being broken down inside the cells, and the specific mechanism on how they are metabolized, could naturally become another focus for further research. Nevertheless, based on these developments and initial validation studies, our work may open new perspectives for developing a treatment for retinal diseases using small hydrophilic cargo.

## 5. Conclusions

This study presents SLN formulation capable of encapsulating small hydrophilic cargo and delivering it to retinal cells *in vitro*. The study highlighted that a gel core could significantly increase the encapsulation efficiency of small hydrophilic cargo inside the SLN. We also observed that the type of hydrophobic polymer used in the composite shell may affect the particle surface charge, a key factor for intravitreal drug delivery system. The physicochemical properties of the DDS developed using RhoB were retained when the neuroprotective cGMP analogue CN03 was used as a cargo. The SLN maintained its particle size below 300 nm after 1 month of storage in PBS. The *in vitro* study demonstrated that the DDS could be uptaken by model retinal cell lines (*i.e*., ARPE-19 and 661W) with different uptake rates based on the particle shell composition and cell type. Equally important, the DDS could release its cargo inside the cells. While the current results are promising for an early-stage formulation development study, more complex *in vivo* studies are needed to demonstrate the clinical relevance of the newly developed DDS.

## Supplementary Materials

The following are available online at www.mdpi.com/xxx/s1, Table S1. Formulation code and component mass dissolved in O-phase for each formulation, Figure S1. Flow cytometry analysis of ARPE-19 cells positive for RhoB after incubation with 200 μg/ml RhoB-SLN.05 or RhoB/SLN.06 at different time point..

## Author Contributions

Study conception and design: E.H., L.H., V.M., M.B., and A.S. Data acquisition: E.H., L.H., S.B., and R.G. Analysis and interpretation of data: E.H. and L.H. Visualization: E.H. and L.H. Original draft preparation: E.H., L.H., S.B., and R.G. Review and editing: V.M., F.P.D., A.S., and N.S. Supervision: V.M., F.P.D., M.B., A.S., and N.S.

## Data Availability Statement

The raw data supporting the conclusions of this article will be made available upon request

## Acknowledgments

We thank Prof Gareth R. Williams from University College London for his assistance in TEM images. We thank Dr. M. Montanari for technical support on flow cytometry.

## References

1. Lechner, J.; O’Leary, O.E.; Stitt, A.W. The pathology associated with diabetic retinopathy. Vision Res. 2017, 139, 7–14, doi:10.1016/j.visres.2017.04.003.

2. Mitchell, P.; Liew, G.; Gopinath, B.; Wong, T.Y. Age-related macular degeneration. Lancet 2018, 392, 1147–1159, doi:10.1016/S0140-6736(18)31550-2.

3. Swaroop, A.; Sieving, P.A. The golden era of ocular disease gene discovery: Race to the finish. Clin. Genet. 2013, 84, 99–101, doi:10.1111/cge.12204.

4. Chakravarthy, U.; Biundo, E.; Saka, R.O.; Fasser, C.; Bourne, R.; Little, J.A. The Economic Impact of Blindness in Europe. Ophthalmic Epidemiol. 2017, 24, 239–247, doi:10.1080/09286586.2017.1281426.

5. Gallego, I.; Villate-Beitia, I.; Martínez-Navarrete, G.; Menéndez, M.; López-Méndez, T.; Soto-Sánchez, C.; Zárate, J.; Puras, G.; Fernández, E.; Pedraz, J.L. Non-viral vectors based on cationic niosomes and minicircle DNA technology enhance gene delivery efficiency for biomedical applications in retinal disorders. Nanomedicine Nanotechnology, Biol. Med. 2019, 17, 308–318, doi:10.1016/j.nano.2018.12.018.

6. Tolone, A.; Belhadj, S.; Rentsch, A.; Schwede, F.; Paquet-Durand, F. The cGMP pathway and inherited photoreceptor degeneration: Targets, compounds, and biomarkers. Genes (Basel). 2019, 10, 1–16, doi:10.3390/genes10060453.

7. Li, Q.; Li, X.; Zhao, C. Strategies to Obtain Encapsulation and Controlled Release of Small Hydrophilic Molecules. Front. Bioeng. Biotechnol. 2020, 8, 1–6, doi:10.3389/fbioe.2020.00437.

8. Himawan, E.; Ekström, P.; Buzgo, M.; Gaillard, P.; Stefánsson, E.; Marigo, V.; Loftsson, T.; Paquet-Durand, F. Drug delivery to retinal photoreceptors. Drug Discov. Today 2019, 24, 1637–1643, doi:10.1016/j.drudis.2019.03.004.

9. Gorantla, S.; Rapalli, V.K.; Waghule, T.; Singh, P.P.; Dubey, S.K.; Saha, R.N.; Singhvi, G. Nanocarriers for ocular drug delivery: Current status and translational opportunity. RSC Adv. 2020, 10, 27835–27855, doi:10.1039/d0ra04971a.

10. Peeters, L.; Sanders, N.N.; Braeckmans, K.; Boussery, K.; Van de Voorde, J.; De Smedt, S.C.; Demeester, J. Vitreous: A Barrier to Nonviral Ocular Gene Therapy. Investig. Opthalmology Vis. Sci. 2005, 46, 3553, doi:10.1167/iovs.05-0165.

11. Xu, Q.; Boylan, N.J.; Suk, J.S.; Wang, Y.Y.; Nance, E.A.; Yang, J.C.; McDonnell, P.J.; Cone, R.A.; Duh, E.J.; Hanes, J. Nanoparticle diffusion in, and microrheology of, the bovine vitreous ex vivo. J. Control. Release 2013, 167, 76–84, doi:10.1016/j.jconrel.2013.01.018.

12. Tavakoli, S.; Kari, O.K.; Turunen, T.; Lajunen, T.; Schmitt, M.; Lehtinen, J.; Tasaka, F.; Parkkila, P.; Ndika, J.; Viitala, T.; et al. Diffusion and Protein Corona Formation of Lipid-Based Nanoparticles in the Vitreous Humor: Profiling and Pharmacokinetic Considerations. Mol. Pharm. 2021, 18, 699–713, doi:10.1021/acs.molpharmaceut.0c00411.

13. del Amo, E.M.; Rimpelä, A.K.; Heikkinen, E.; Kari, O.K.; Ramsay, E.; Lajunen, T.; Schmitt, M.; Pelkonen, L.; Bhattacharya, M.; Richardson, D.; et al. Pharmacokinetic aspects of retinal drug delivery. Prog. Retin. Eye Res. 2017, 57, 134–185, doi:10.1016/j.preteyeres.2016.12.001.

14. Apaolaza, P.S.; del Pozo-Rodríguez, A.; Solinís, M.A.; Rodríguez, J.M.; Friedrich, U.; Torrecilla, J.; Weber, B.H.F.; Rodríguez-Gascón, A. Structural recovery of the retina in a retinoschisin-deficient mouse after gene replacement therapy by solid lipid nanoparticles. Biomaterials 2016, 90, 40–49, doi:10.1016/j.biomaterials.2016.03.004.

15. Bisht, R.; Mandal, A.; Jaiswal, J.K.; Rupenthal, I.D. Nanocarrier mediated retinal drug delivery: overcoming ocular barriers to treat posterior eye diseases. Wiley Interdiscip. Rev. Nanomedicine Nanobiotechnology 2018, 10, 1–21, doi:10.1002/wnan.1473.

16. Zariwala, M.G.; Bendre, H.; Markiv, A.; Farnaud, S.; Renshaw, D.; Taylor, K.M.G.; Somavarapu, S. Hydrophobically modified chitosan nanoliposomes for intestinal drug delivery. Int. J. Nanomedicine 2018, 13, 5837–5848, doi:10.2147/IJN.S166901.

17. Khalkhali, M.; Mohammadinejad, S.; Khoeini, F.; Rostamizadeh, K. Vesicle-like structure of lipid-based nanoparticles as drug delivery system revealed by molecular dynamics simulations. Int. J. Pharm. 2019, 559, 173–181, doi:10.1016/j.ijpharm.2019.01.036.

18. Severino, P.; Pinho, S.C.; Souto, E.B.; Santana, M.H.A. Polymorphism, crystallinity and hydrophilic–lipophilic balance of stearic acid and stearic acid–capric/caprylic triglyceride matrices for production of stable nanoparticles. Colloids Surfaces B Biointerfaces 2011, 86, 125–130, doi:10.1016/j.colsurfb.2011.03.029.

19. Yang, R.; Gao, R.C.; Cai, C.F.; Xu, H.; Li, F.; He, H.B.; Tang, X. Preparation of gel-core-solid lipid nanoparticle: A novel way to improve the encapsulation of protein and peptide. Chem. Pharm. Bull. 2010, 58, 1195–1202, doi:10.1248/cpb.58.1195.

20. Chen, C.; Zhu, X.; Dou, Y.; Xu, J.; Zhang, J.; Fan, T.; Du, J.; Liu, K.; Deng, Y.; Zhao, L.; et al. Exendin-4 loaded nanoparticles with a lipid shell and aqueous core containing micelles for enhanced intestinal absorption. J. Biomed. Nanotechnol. 2015, 11, 865–876, doi:10.1166/jbn.2015.1971.

21. Xu, Y.; Zheng, Y.; Wu, L.; Zhu, X.; Zhang, Z.; Huang, Y. Novel Solid Lipid Nanoparticle with Endosomal Escape Function for Oral Delivery of Insulin. ACS Appl. Mater. Interfaces 2018, 10, 9315–9324, doi:10.1021/acsami.8b00507.

22. Martens, T.F.; Remaut, K.; Demeester, J.; De Smedt, S.C.; Braeckmans, K. Intracellular delivery of nanomaterials: How to catch endosomal escape in the act. Nano Today 2014, 9, 344–364, doi:10.1016/j.nantod.2014.04.011.

23. Dunn, K.C.; Aotaki-Keen, A.E.; Putkey, F.R.; Hjelmeland, L.M. ARPE-19, A Human Retinal Pigment Epithelial Cell Line with Differentiated Properties. Exp. Eye Res. 1996, 62, 155–170, doi:10.1006/exer.1996.0020.

24. Tan, E.; Ding, X.-Q.; Saadi, A.; Agarwal, N.; Naash, M.I.; Al-Ubaidi, M.R. Expression of Cone-Photoreceptor–Specific Antigens in a Cell Line Derived from Retinal Tumors in Transgenic Mice. Investig. Opthalmology Vis. Sci. 2004, 45, 764, doi:10.1167/iovs.03-1114.

25. Chen, C.; Fan, T.; Jin, Y.; Zhou, Z.; Yang, Y.; Zhu, X.; Zhang, Z.R.; Zhang, Q.; Huang, Y. Orally delivered salmon calcitonin-loaded solid lipid nanoparticles prepared by micelle-double emulsion method via the combined use of different solid lipids. Nanomedicine 2013, 8, 1085–1100, doi:10.2217/nnm.12.141.

26. Vighi, E.; Trifunovic, D.; Veiga-Crespo, P.; Rentsch, A.; Hoffmann, D.; Sahaboglu, A.; Strasser, T.; Kulkarni, M.; Bertolotti, E.; Van Den Heuvel, A.; et al. Combination of cGMP analogue and drug delivery system provides functional protection in hereditary retinal degeneration. Proc. Natl. Acad. Sci. U. S. A. 2018, 115, E2997–E3006, doi:10.1073/pnas.1718792115.

27. Firdessa, R.; Oelschlaeger, T.A.; Moll, H. Identification of multiple cellular uptake pathways of polystyrene nanoparticles and factors affecting the uptake: Relevance for drug delivery systems. Eur. J. Cell Biol. 2014, 93, 323–337, doi:10.1016/j.ejcb.2014.08.001.

28. Yang, S.T.; Zaitseva, E.; Chernomordik, L. V.; Melikov, K. Cell-penetrating peptide induces leaky fusion of liposomes containing late endosome-specific anionic lipid. Biophys. J. 2010, 99, 2525–2533, doi:10.1016/j.bpj.2010.08.029.

29. Foroozandeh, P.; Aziz, A.A. Insight into Cellular Uptake and Intracellular Trafficking of Nanoparticles. Nanoscale Res. Lett. 2018, 13, 339, doi:10.1186/s11671-018-2728-6.

30. Kong, B.; Seog, J.H.; Graham, L.M.; Lee, S.B. Experimental considerations on the cytotoxicity of nanoparticles. Nanomedicine 2011, 6, 929–941, doi:10.2217/nnm.11.77.

31. He, C.; Hu, Y.; Yin, L.; Tang, C.; Yin, C. Effects of particle size and surface charge on cellular uptake and biodistribution of polymeric nanoparticles. Biomaterials 2010, 31, 3657–3666, doi:10.1016/j.biomaterials.2010.01.065.

32. Kumari, S.; Mg, S.; Mayor, S. Endocytosis unplugged: multiple ways to enter the cell. Cell Res. 2010, 20, 256–275, doi:10.1038/cr.2010.19.

33. Bertolotti, E.; Neri, A.; Camparini, M.; Macaluso, C.; Marigo, V. Stem cells as source for retinal pigment epithelium transplantation. Prog. Retin. Eye Res. 2014, 42, 130–144, doi:10.1016/j.preteyeres.2014.06.002.

34. Behzadi, S.; Serpooshan, V.; Tao, W.; Hamaly, M.A.; Alkawareek, M.Y.; Dreaden, E.C.; Brown, D.; Alkilany, A.M.; Farokhzad, O.C.; Mahmoudi, M. Cellular uptake of nanoparticles: journey inside the cell. Chem. Soc. Rev. 2017, 46, 4218–4244, doi:10.1039/C6CS00636A.

35. Palocci, C.; Valletta, A.; Chronopoulou, L.; Donati, L.; Bramosanti, M.; Brasili, E.; Baldan, B.; Pasqua, G. Endocytic pathways involved in PLGA nanoparticle uptake by grapevine cells and role of cell wall and membrane in size selection. Plant Cell Rep. 2017, 36, 1917–1928, doi:10.1007/s00299-017-2206-0.

36. Arana, L.; Bayón-Cordero, L.; Sarasola, L.; Berasategi, M.; Ruiz, S.; Alkorta, I. Solid Lipid Nanoparticles Surface Modification Modulates Cell Internalization and Improves Chemotoxic Treatment in an Oral Carcinoma Cell Line. Nanomaterials 2019, 9, 464, doi:10.3390/nano9030464.

37. Belhadj, S.; Tolone, A.; Christensen, G.; Das, S.; Chen, Y.; Paquet-Durand, F. Long-Term, Serum-Free Cultivation of Organotypic Mouse Retina Explants with Intact Retinal Pigment Epithelium. J. Vis. Exp. 2020, doi:10.3791/61868.

