## Supplemental Table S1 and Supplemental Figure S1 for "Efficient Hydrophilic Small Molecule Delivery To Retinal Cell Lines Using Solid Lipid Nanoparticle Containing Gel Core"

Table of Contents

Table S1
Figure S1

Table S1. Formulation code and component mass dissolved in O-phase for each formulation.

| **Drug Delivery System (DDS)** | **Tripalmitin (GTP)** | **Soy-Bean Lecithin (LCT)** | **Stearic Acid (SA)** | **Poly-Ɛ-caprolactone (PCL)** | **50/50 DL-lactide/glycolide (PLGA)** | **Core** |
|---|---|---|---|---|---|---|
| SLN.01 | 15 | 15 | 1 | - | - | Aqueous |
| SLN.02 | 15 | 15 | 1 | 10 | - | Aqueous |
| SLN.03 | 15 | 15 | 1 | - | 10 | Aqueous |
| SLN.04 | 15 | 15 | 1 | - | - | Aqueous |
| SLN.05 | 15 | 15 | 1 | 10 | - | Hydrogel |
| SLN.06 | 15 | 15 | 1 | - | 10 | Hydrogel |

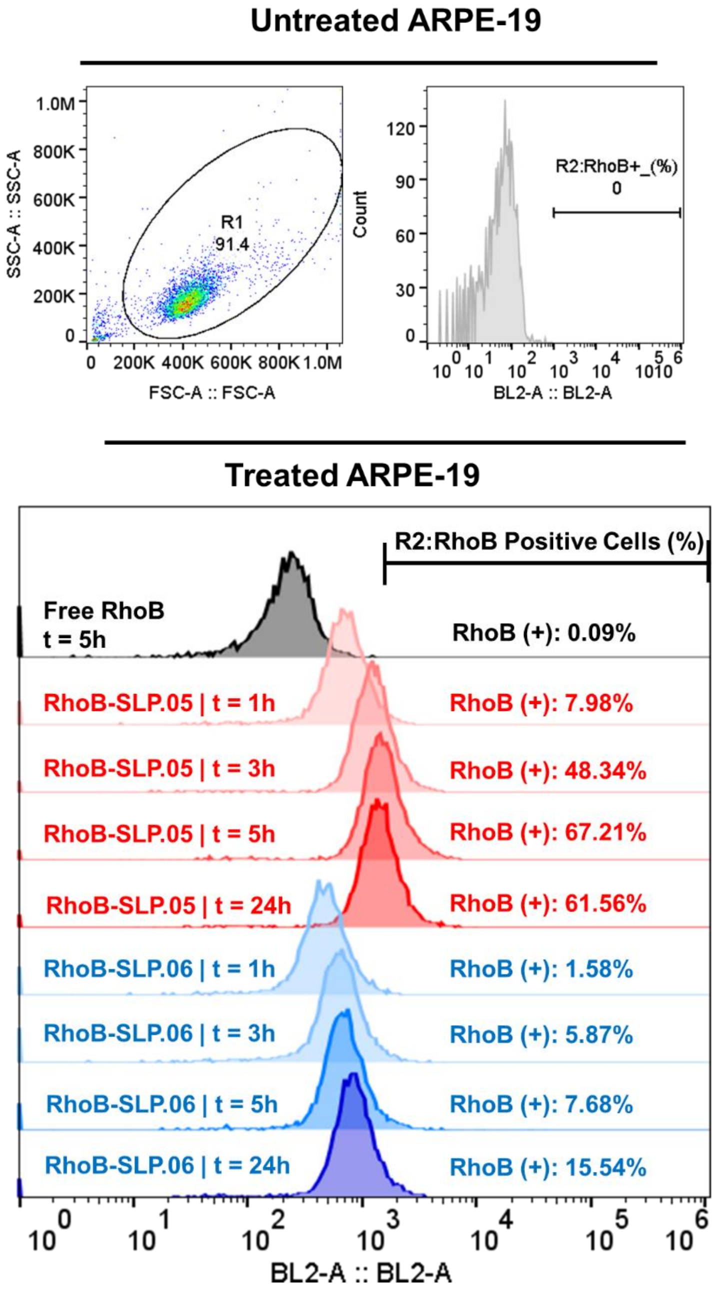


**Supplementary Figure S1. Flow cytometry analysis of ARPE-19 cells positive for RhoB after incubation with 200 µg/ml RhoB-SLN.05 or RhoB/SLN.06 at different time point.**

**(A)** Gating on ARPE-19 cells to determine RhoB positive cells.

**(B)** Histogram overlay of RhoB relative fluorescence intensity and percentage of RhoB positive cell (RhoB+). R1: gating for selecting cells population; R2: gating to select RhoB+ cells based on blue laser (BL2-A) detector; FSC-A: forward scattering channel; SSC-A: side scattering chanel.
